# ATR inhibition enhances 5-fluorouracil sensitivity independent of non-homologous end-joining and homologous recombination repair pathway

**DOI:** 10.1101/2020.04.20.051318

**Authors:** Soichiro S. Ito, Yosuke Nakagawa, Masaya Matsubayashi, Yoshihiko M. Sakaguchi, Shinko Kobashigawa, Takeshi K. Matsui, Hitoki Nanaura, Mari Nakanishi, Fumika Kitayoshi, Sotaro Kikuchi, Atsuhisa Kajihara, Shigehiro Tamaki, Kazuma Sugie, Genro Kashino, Akihisa Takahashi, Masatoshi Hasegawa, Eiichiro Mori, Tadaaki Kirita

**Affiliations:** Department of Oral and Maxillofacial Surgery, Nara Medical University, Kashihara, Nara 634-8522, Japan; Department of Future Basic Medicine, Nara Medical University, Kashihara, Nara 634-8521, Japan; Department of Neurology, Nara Medical University, Kashihara, Nara 634-8522, Japan; Radioisotope Research Center, Nara Medical University, Kashihara, Nara 634-8521, Japan; Gunma University Heavy Ion Medical Center, Maebashi, Gunma 371-8511, Japan; Department of Radiation Oncology, Nara Medical University, Kashihara, Nara 634-8522, Japan

**Keywords:** ATR, 5-fluorouracil, cell cycle checkpoint, DNA double-strand breaks, homologous recombination, BRCA2

## Abstract

The anticancer agent, 5-fluorouracil (5-FU), is typically applied in the treatment of various types of cancers because of its properties. Thought to be an inhibitor of the enzyme thymidylate synthase which plays a role in nucleotide synthesis, 5-FU has been found to induce single- and double-strand DNA breaks. The activation of ATR occurs as a reaction to UV- and chemotherapeutic drug-induced replication stress. In this study, we examined the effect of ATR inhibition on 5-FU sensitivity. Using western blotting, we found that 5-FU treatment led to the phosphorylation of ATR. Surviving fractions were remarkably decreased in 5-FU with ATR inhibitor (ATRi) compared to 5-FU with other major DNA repair kinases inhibitors. ATR inhibition enhanced induction of DNA double-strand breaks and apoptosis in 5-FU-treated cells. Using gene expression analysis, we found that 5-FU could induce the activation of intra-S checkpoint. Surprisingly, *BRCA2*-deficient cells were sensitive to 5-FU in the presence of ATRi. In addition, ATR inhibition enhanced the efficacy of 5-FU treatment, independent of non-homologous end-joining and homologous recombination repair pathways. Findings from the present study suggest ATR as a potential therapeutic target for 5-FU chemotherapy.

---

One of many anticancer agents available, 5-fluorouracil (5-FU) has nonetheless become the drug-of-choice in the treatment of various solid tumors because of its properties. It converts to a number of active metabolites (such as fluorouridine triphosphate {FUTP}, fluorodeoxyuridine triphosphate {FdUTP}, and fluorodeoxyuridine monophosphate {FdUMP}) that disrupt RNA and DNA metabolism, and inhibit thymidylate synthase (TS) (1,2). Specifically, global RNA metabolism is compromised when the following conditions take place: 1) RNA absorbs the 5-FU converted active metabolite, FUTP, in place of uridine triphosphate (UTP); and 2) FdUTP, instead of deoxythymidine triphosphate (dTTP), is absorbed by DNA (2,3). A third process occurs and results in DNA damage. The absorption of FdUMP contributes to an inhibition of TS via the formation of a ternary covalent complex that consists of TS-FdUMP-5, 10-methylenetetrahydrofolate. It also blocks cells from the synthesis of dTMP from dUMP, causing cellular dUTP increases at the expense of dTTP. This leads to a significant misincorporation of dUTP or FdUTP during replication, and in particular, that of DNA.

Despite consistent observations of DNA damage as one of 5-FU mediated tumor cell killings (4,5), the exact mechanism behind the processing and contribution of misincorporated dUTP and FdUTP to cytotoxicity has yet to be fully elucidated. Studies suggest that base excision repair (BER) enzymes and mismatch repair (MMR) identify misincorporated FdUTP and dUTP for excision from DNA (5–7). The BER enzyme uracil-DNA-glycosylase initiates repair of DNA through the elimination of uracil or 5-FU from DNA (8). However, this repair mechanism is rendered ineffective given high FdUTP/dTTP ratios, and only serves to trigger the incorporation of additional false nucleotides (4). The MMR system plays an equally significant role in the correction of such replication errors via the nicks and gaps in single-strand DNA (ssDNA) produced by the FdUTP and dUTP in both BER and MMR (4). These nicks and gaps trigger the initial activation of ATR-checkpoint kinase 1 (Chk1) pathways, and activated Chk1 molecules, in turn, stop DNA replication. During these processes, unstable conformations in the DNA structure are induced by the coating of stalled replication fork complexes with replication protein A. The presence of excessive single-strand breaks (SSBs) at stalled replication forks subsequently induces DNA double-strand breaks (DSBs) in 5-FU treated cells (9,10).

DNA damage response (DDR) involves the activation of a signaling network that provides time for DNA repair and triggers apoptosis when extensive damage occurs. It effectively mediates cell cycle arrest and is triggered by the activation of protein kinases, ataxia telangiectasia mutated (ATM), ATM and Rad3-related (ATR), and DNA-dependent protein kinase catalytic subunit (DNA-PKcs), which is one of three members of the phosphoinositide-3-kinase-like protein kinase (PIKK) family. Cell cycle progression in G1, S, or G2 phase is delayed as these kinases recruit repair machinery to damaged DNA sites via the activation of effector checkpoints (11). Through p21/CIP1/WAF1 upregulation, p53 mediates G1 arrest and, where extensive DNA damage is detected, triggers apoptosis (12). Nonetheless, as a majority of cancer cells display loss of p53 function and its regulatory pathways, it is evident that chemotherapy-induced DNA damage is unable to halt G1 phase or promote apoptosis. Cells rely solely upon S and G2/M checkpoints for the arrest of cell cycle and facilitation of DNA repair post genotoxic exposure and prior to mitosis. ATR/Chk1 kinases have been found to be implicated in the regulation of post genotoxic stress cell cycle arrest, impediment of subsequent replication origin firing during S phase, and involvement in the intra-S and G2/M checkpoints (13–16).

Homologous recombination (HR) repair is one of the major DSBs repair pathways (17,18) which operates primarily via intact sister chromatids during late S and G2 phases, but not at G1 phase (19,20). BRCA2, Rad52, Rad54 and Rad51 paralogs such as Rad51C-XRCC3 and Rad51B-Rad51C-Rad51D-XRCC2 are some of the proteins involved in vertebrate cells HR (21). BRCA2, an upstream protein, has been shown to regulate Rad51 activity (22). Mutations in the BRCA2 gene have consistently been found in hereditary breast (23) and ovarian cancers (24).

We previously demonstrated that BRCA2, a major component of HR repair pathway, plays a crucial role in protecting cells from cell death and in the repair of DNA damage induced by 5-FU (3). However, the manner in which cells detect and respond to DNA damage induced by 5-FU remains unclear. ATR is activated in response to replication stress induced by DNA damaging reagents, and acts upon the upstream of BRCA2-dependent repair pathway (25–28). ATR is one of the principal kinases of the DNA damage response, in addition to ATM and DNA-PK. In the present study, we sought to characterize the role of ATR in response to 5-FU and examine the effect of an ATR inhibitor for cancer treatment with 5-FU.

## Results

### ATR inhibition sensitized 5-FU-treated cells

To examine the activation of ATR and other major DDR kinases by 5-FU treatment, we verified the phosphorylation of ATR, ATM, DNA-PKcs and Chk1 by western blotting in SAS cells. The phosphorylation of ATR at Thr-1989 (an autophosphorylation site; 29) was induced by 5-FU in a dose-dependent manner (Fig. 1). Our observations suggest that 5-FU treatment activated ATR. On the other hand, the phosphorylation of ATM at Ser-1981 induced by 5-FU reached peak levels at 20 μM. More phosphorylation of DNA-PKcs at Ser-2056 induced by X-rays (20 Gy) irradiation was detected than that induced by 5-FU. Like the ATR phosphorylation, Chk1 phosphorylation at Ser-345 induced by 5-FU was detected in a dose-dependent manner (Fig. 1).

**Figure 1.**
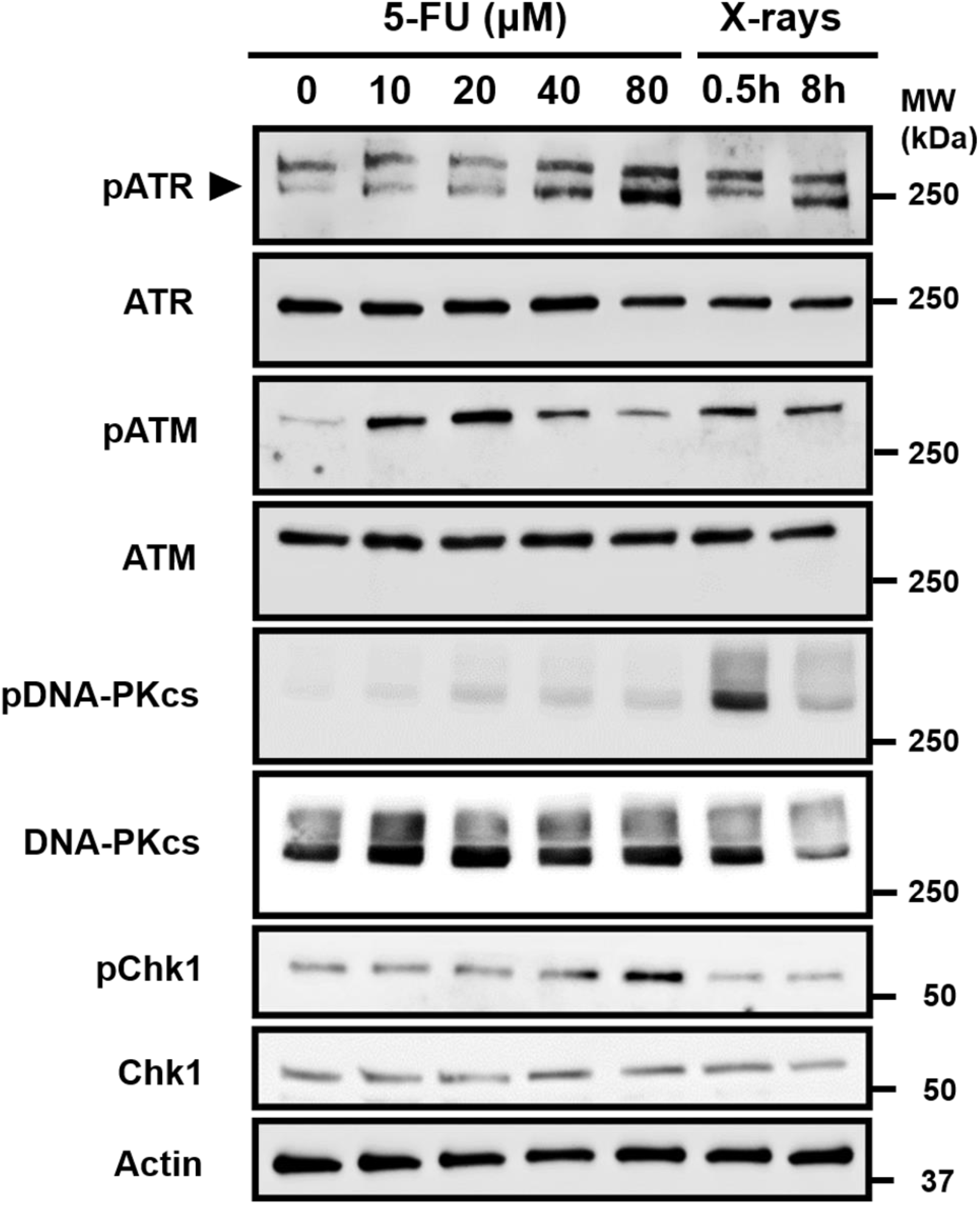
Phosphorylation of three members of the PIKK family and Chk1. Phosphorylation of ATR, ATM, DNA-PKcs and Chk1 by 5-FU treatment for 24 h. Cells were collected immediately after treatment. Irradiated cells were collected at 0.5 h and 8 h after X-rays-irradiation (20 Gy). Arrow indicates phosphorylated form of ATR. Upper bands are non-specific. All experiments were replicated 3 times.

To check whether the concentration of a specific inhibitor against ATR (ATRi) alone does not exceed IC_50_ in our experiments, we analyzed cell survival. Surviving fraction showed slight decrease under 10 μM ATRi (Fig. S1A and B), and IC_50_ value was approximately 15 μM in Chinese hamster lung fibroblasts (Fig. S1A) and 5 μM in SAS cells (Fig. S1B). Although 3 μM ATRi alone did not have cell lethality, it synergistically enhanced the cytotoxicity of 5-FU treatment (Fig. 2A).

**Figure 2.**
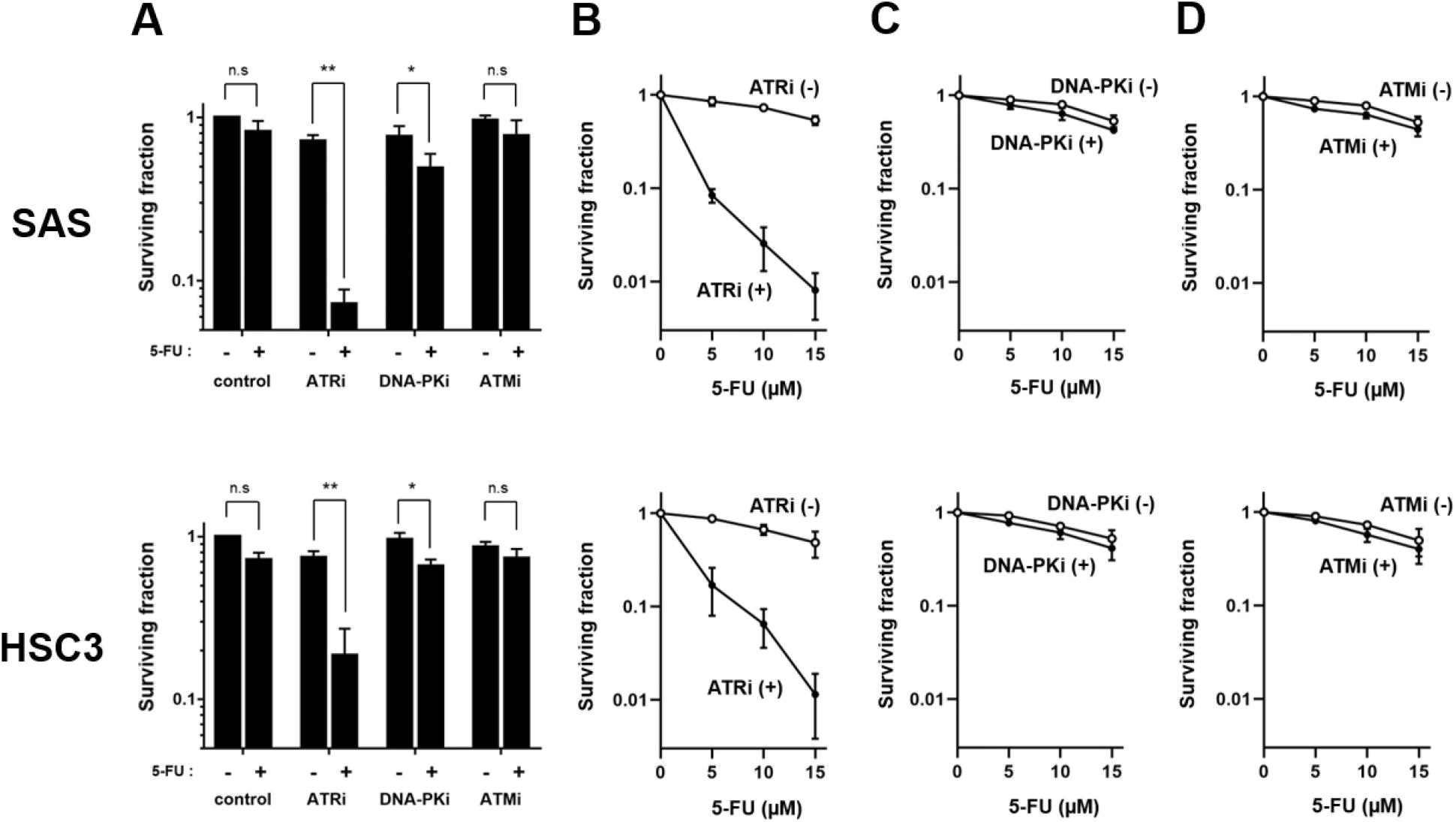
Sensitivity of three members of the PIKK family inhibition against 5-FU treated SAS and HSC3 cells. (A) ATRi, DNA-PKi and ATMi were used at 3 μM respectively. 5-FU was used at 5 μM. Surviving fraction was significantly decreased in 5-FU treatment combined with ATRi compared to treatments combined with either DNA-PKi or ATMi. (B, C, D) Surviving fraction in 5-FU only treatment (open circles) was compared against 5-FU treatment combined with 3 μM ATRi (B)/DNA-PKi (C)/ATMi (D) (filled circles) in SAS (upper column) and HSC3 (lower column). All experiments were replicated 3 times. The values obtained were described as means ± SD. Data were compared statistically using the two-tailed Student’s *t*-test. *p-*value; *, ** and *** represent *p* < 0.05, *p* < 0.01 and *p* < 0.001, respectively.

To verify the significance of ATRi compared to other major DDR kinase inhibitors against DNA-PK (DNA-PKi) and ATM (ATMi), SAS and HSC3 cells were exposed to 5 μM 5-FU, and 3 μM ATRi/DNA-PKi/ATMi for 24 h, and subsequently measured using a standard colony forming assay. The number of surviving fractions were significantly lower in 5-FU treatment combined with ATRi, compared to 5-FU treatment combined with DNA-PKi or ATMi (Fig. 2A). We confirmed cell viability by changing the concentration of 5-FU. Surviving fraction was remarkably decreased in the 5-FU treatment combined with ATRi compared to 5-FU alone in both SAS and HSC3 cells (Fig. 2B). The number of surviving fractions were lower in 5-FU treatment combined with ATRi, compared to 5-FU treatment combined with DNA-PKi or ATMi (Fig. 2C and D). Surviving fraction decreases in 5-FU combined with ATRi are assumed to occur in a *p53*-independent manner (Fig. S1C). These findings suggest that ATRi is more effective in 5-FU treatment than DNA-PKi or ATMi. Figures 3A and B demonstrate that ATRi suppressed the phosphorylation of ATR and Chk1 induced by 5-FU, suggesting that ATRi blocks both ATR autophosphorylation and Chk1 phosphorylation.

**Figure 3.**
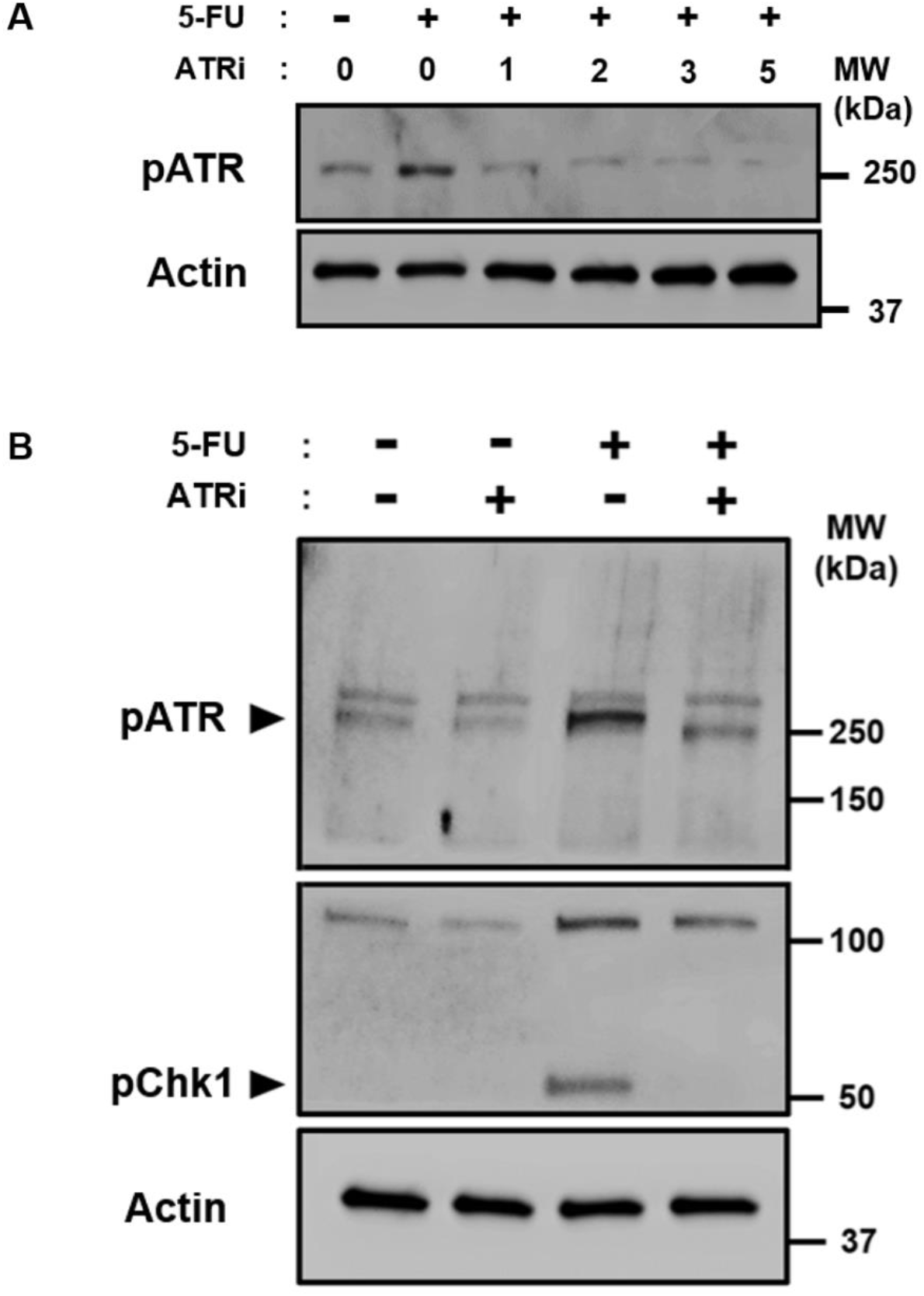
The phosphorylation of ATR and Chk1 induced by 5-FU treatment was suppressed by ATR inhibitor VE-821. (A) Expression analysis by western blotting of ATR. ATRi was used at 1 μM, 2 μM, 3 μM, and 5 μM. 5-FU were used at 10 μM. Cells were simultaneously treated by both chemicals for 24 h. (B) Expression analysis by western blotting of ATR and Chk1. ATRi was used at 3 μM. 5-FU were used at 10 μM. Cells were simultaneously treated by both chemicals for 24 h. Cells were collected immediately after treatment. ATR and its major downstream effector, Chk1, were phosphorylated by 5-FU treatment. In addition, these phosphorylations were suppressed by ATRi. Arrow indicates phosphorylated form of ATR and Chk1. Upper bands are non-specific. All experiments were replicated 3 times.

### ATR inhibition enhanced induction of DSBs in 5-FU-treated cells

To confirm the degree of DSB induction, we performed comet assays under neutral conditions. SAS cells were treated with 10 μM 5-FU and/or 3 μM ATRi for 12 h. Tail moments of 5-FU treatment combined with ATRi were significantly increased (Fig. 4A, B and S2). In addition, we performed another method for detecting DSBs. γH2AX immunocytochemical staining is a sensitive method by which DSBs can be detected (30). This was used to examine the presence of H2AX phosphorylation induced by 5-FU and ATRi. Figure 5A depicts a typical H2AX phosphorylation in SAS cells after 12-h treatment with 10 μM 5-FU and 3 μM ATRi combined. To quantify the optical intensity of H2AX phosphorylation using flow cytometry, cells were exposed to 10 μM 5-FU with or without 3 μM ATRi for 6 h and 12 h (Fig. 5B). The intensity of H2AX phosphorylation treated with 5-FU alone for 6 h was 103.7 ± 0.6, and for 12 h was 119.1 ± 0.7, whereas the intensity after a 5-FU combined with ATRi treatment for 6 h was 105.4 ± 0.3, and for 12 h was 162.1 ± 1.8 in SAS cells (Fig. 5C). The intensity of H2AX phosphorylation treated with 5-FU alone for 6 h was 110.6 ± 0.8, and for 12 h was 237.9 ± 2.3, and post 5-FU combined with ATRi treatment for 6 h was 118.7 ± 0.6, and for 12 h was 364.3 ± 3.8 in HSC3 cells (Fig. 5C). Our findings suggest that ATR inhibition leads to a less efficient repair of 5-FU-induced DNA damage.

**Figure 4.**
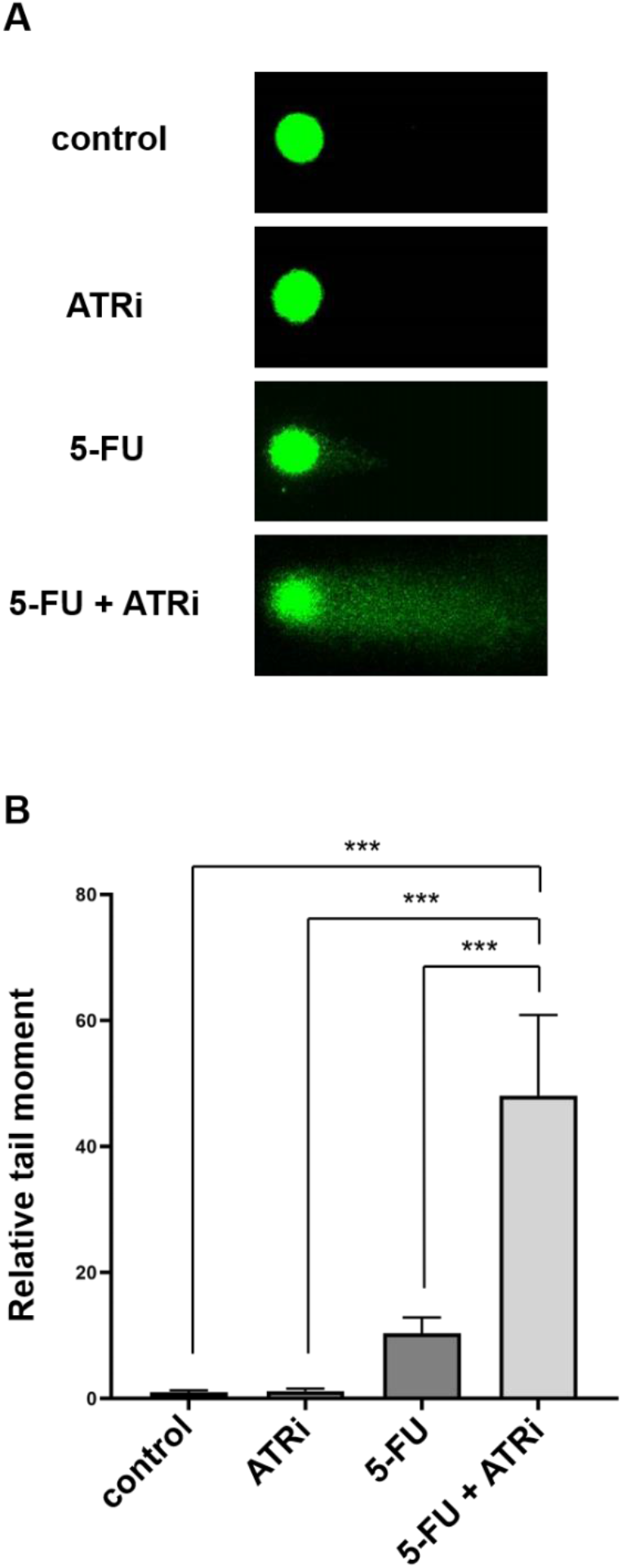
DSBs analysis by neutral comet assay. (A) Typical images of comet assays in SAS. Cells were exposed to 10 μM 5-FU and/or 3 μM ATRi treatment for 12 h. (B) The tail moments of more than 50 cells were quantified, respectively. All experiments were replicated 3 times. The values obtained were described as means ± SD. Data were compared statistically using the two-tailed Student’s *t*-test. *p-*value; *, ** and *** represent *p* < 0.05, *p* < 0.01 and *p* < 0.001, respectively.

**Figure 5.**
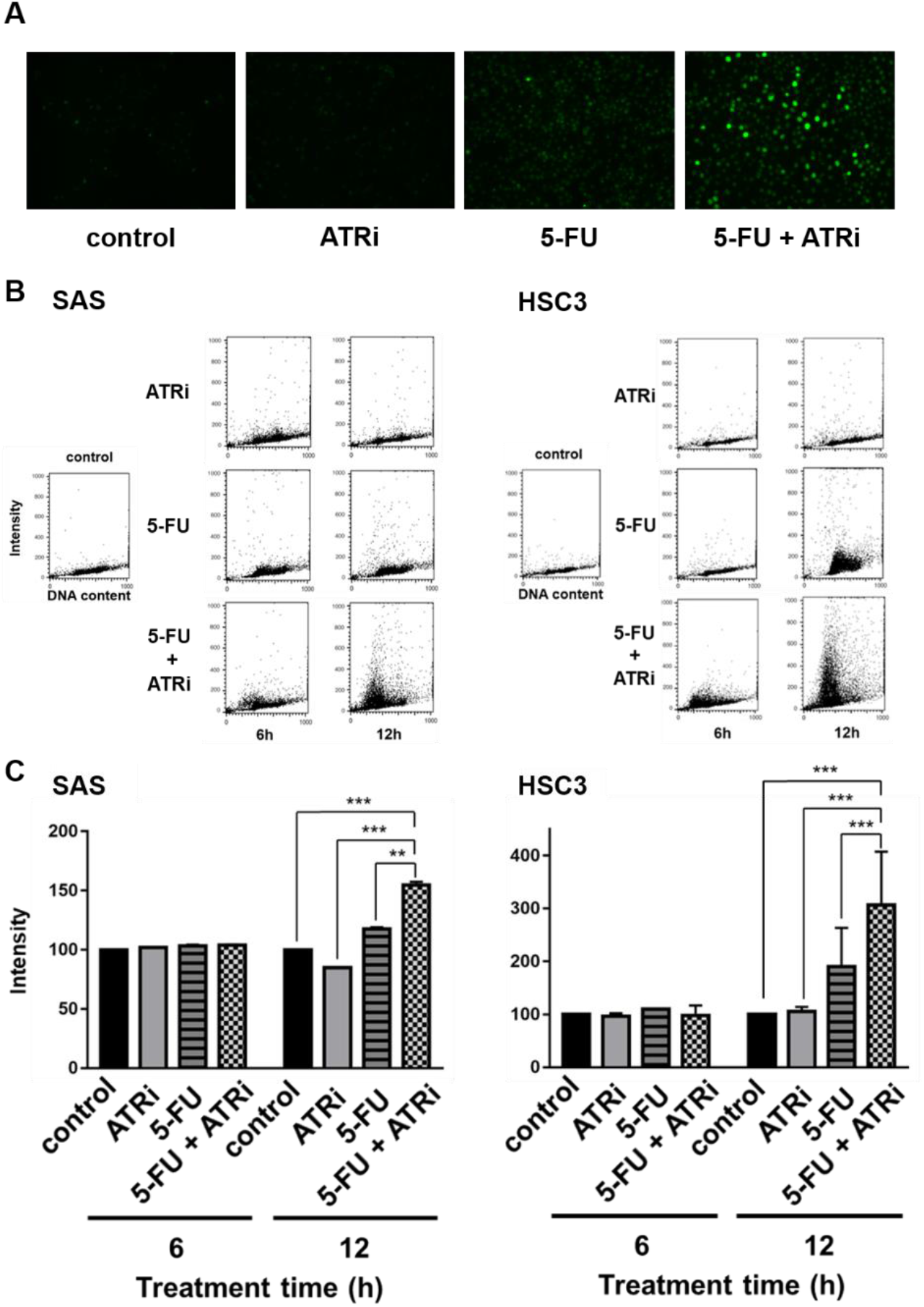
Histone H2AX phosphorylation analysis. (A) Phosphorylation of histone H2AX was detected in SAS. Cells were exposed to 10 μM 5-FU and/or 3 μM ATRi treatment for 12 h. (B) Phosphorylation of histone H2AX was measured using flow cytometry in SAS and HSC3. Cells were exposed to 10 μM 5-FU and/or 3 μM ATRi treatment for 6 h and 12 h. (C) Its intensity was schematized as illustrated in SAS and HSC3. All experiments were replicated 3 times. The values obtained were described as means ± SD. Data were compared statistically using the two-tailed Student’s *t*-test. *p-*value; *, ** and *** represent *p* < 0.05, *p* < 0.01 and *p* < 0.001, respectively.

### ATR inhibition enhanced apoptosis induced by 5-FU

To assess the apoptosis induction, cells were detected and quantified with Hoechst33342 staining assay (Fig. 6A). The fraction of apoptosis by 3 μM ATRi alone, 10 μM 5-FU alone, and 10 μM 5-FU combined with 3 μM ATRi for 12 h was 12.9 ± 5.3%, 15.6 ± 6.9%, and 40.5 ± 5.1% respectively, and 11.3 ± 2.0%, 30.6 ± 5.8%, and 59.8 ± 15.5% respectively, for 24 h in SAS cells (Fig. 6B). Counterparts for 12 h was 8.6 ± 3.8%, 15.5 ± 8.5%, and 30.0 ± 16.6% respectively, and 15.6 ± 6.9%, 46.0 ± 11.0%, and 66.4 ± 11.5% respectively, for 24 h in HSC3 cells (Fig. 6B). Apoptotic bodies appeared at a higher frequency in cells given 5-FU treatment combined with ATRi both in SAS and HSC3 cells (Fig. 6A and B).

**Figure 6.**
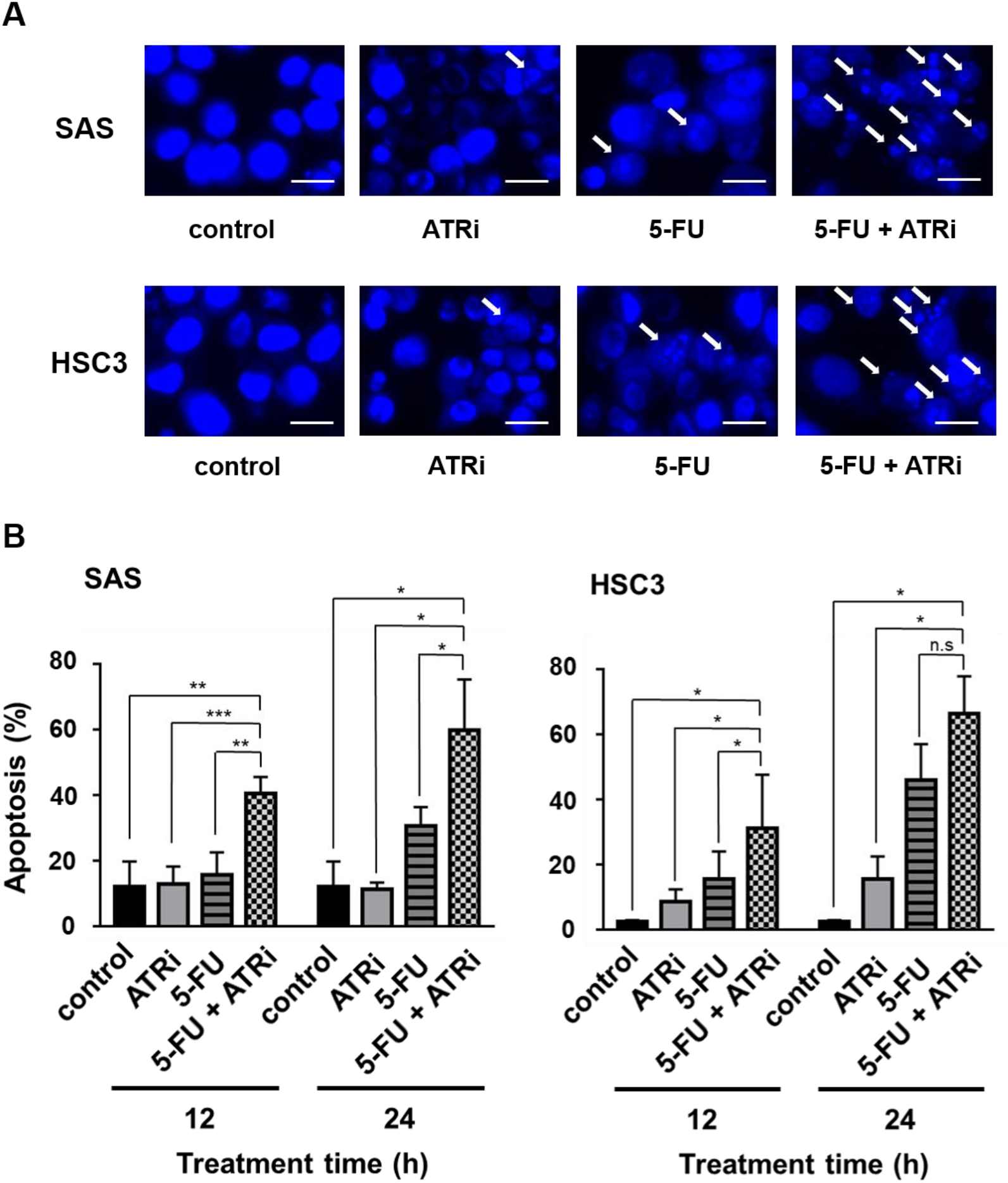
Apoptosis analysis by Hoechst staining. (A) For morphological assessment, apoptotic cells were detected and quantified with Hoechst33342 staining assay in SAS and HSC3. (B) Cells were exposed 10 μM 5-FU and/or 3 μM ATRi for 12 h/24 h. The fraction of apoptosis was schematized as illustrated SAS and HSC3. Bars = 20μm. All experiments were replicated 3 times. The values obtained were described as means ± SD. Data were compared statistically using the two-tailed Student’s *t*-test. *p-*value; *, ** and *** represent *p* < 0.05, *p* < 0.01 and *p* < 0.001, respectively.

To analyze cell cycle profile after 5-FU treatment, we examined the cell cycle distribution. When DNA fragmentation occurred, the position of apoptotic cells was shifted to lower DNA content values, and sub-G1 population was detected far left from G1-peak (31,32). Cells were exposed to 10 μM 5-FU and/or 3 μM ATRi for 8 h/16 h/24 h (Fig. 7A). Treatment with 5-FU alone for 8 h, 16 h, and 24 h, fraction of sub-G1 were 3.3 ± 0.3%, 14.8 ± 0.2%, and 38.1 ± 2.8%, and treatment with 5-FU and ATRi for 8 h, 16 h, and 24 h were 8.8 ± 0.1%, 49.6 ± 3.8%, and 68.1 ± 1.0% in SAS cells (Fig. 7B). Counterparts were 4.3 ± 0.1%, 12.2 ± 0.2%, and 15.1 ± 0.3%, 18.8 ± 0.6%, 46.0 ± 1.7%, and 68.2 ± 1.4% in HSC3 cells (Fig. 7B). Fraction of sub-G1 was remarkably increased in 5-FU treatment combined with ATRi compared to 5-FU treatment alone.

**Figure 7.**
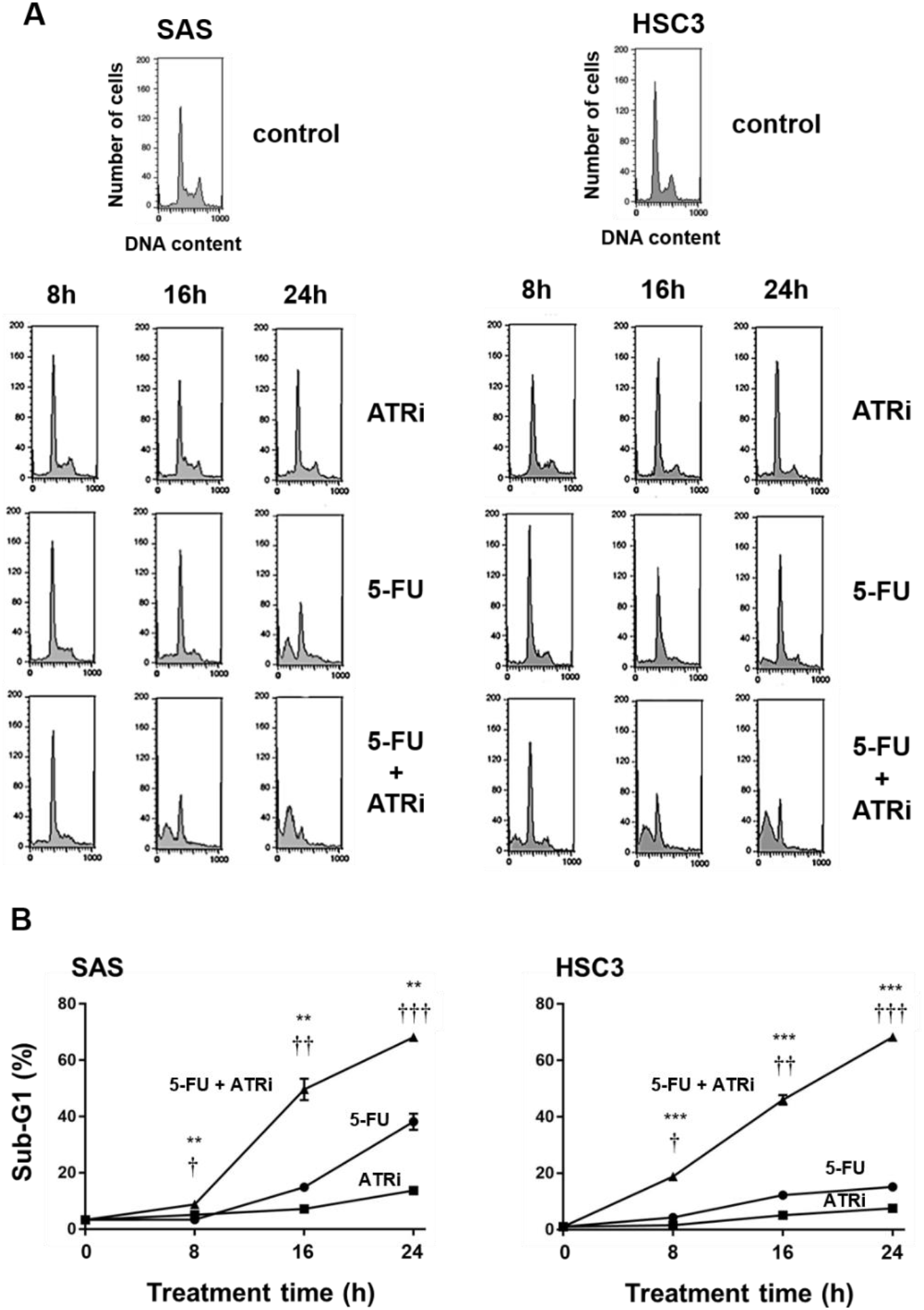
Apoptosis analysis by sub-G1 accumulation. (A) Results of cell cycle analysis after 10 μM 5-FU and/or 3 μM ATRi treatment for 8 h/16 h/24 h in SAS and HSC3. (B) Sub-G1 were schematized as illustrated SAS and HSC3. All experiments were replicated 3 times. The values obtained were described as means ± SD. Data were compared statistically using the two-tailed Student’s *t*-test. *p-*value; */ †, **/ †† and ***/ ††† represent *p* < 0.05, *p* < 0.01 and *p* < 0.001, respectively.

### 5-FU induced cell cycle arrest between S phase and mitotic phase

5-FU treatment caused cell cycle arrest in S phase at 8 and 16 h (Fig. 7A). To investigate the nature of the cell cycle arrest involved in the response to 5-FU treatment, we analyzed genome-wide mRNA by bulk RNA sequencing. The 32 genes that regulated significantly were detected by differentially expressed gene (DEG) analysis both in SAS and HSC3 (Fig. 8A, B, S3A, B, C and Table. S1). We then screened 9 genes associated with cell cycle from the 32 genes by Gene Ontology (GO) analysis. The expression of 4 genes, *Cyclin E1* (*CCNE1*), *Cyclin E2* (*CCNE2*), *Cyclin dependent kinase inhibitor 1A* (*CDKN1A*) and *Thioredoxin interacting protein* (*TXNIP*) were upregulated, and 5 genes, *Cyclin B1* (*CCNB1*), *Cyclin dependent kinase inhibitor 3* (*CDKN3*), *Cell division cycle 20* (*CDC20*), *Aurora kinase A* (*AURKA*) and *Proline and serine rich coiled-coil 1* (*PSRC1*) were downregulated after 10 μM 5-FU treatment for 16 h (Fig. 8B). Subsequently, we quantified these gene expressions by qRT-PCR in each cell line, and similar tendencies were indicated in all the genes listed above (Fig. 9A and B). Our data supports the idea that 5-FU treatment leads to cell cycle arrest between S phase and mitotic phase, particularly in S phase.

**Figure 8.**
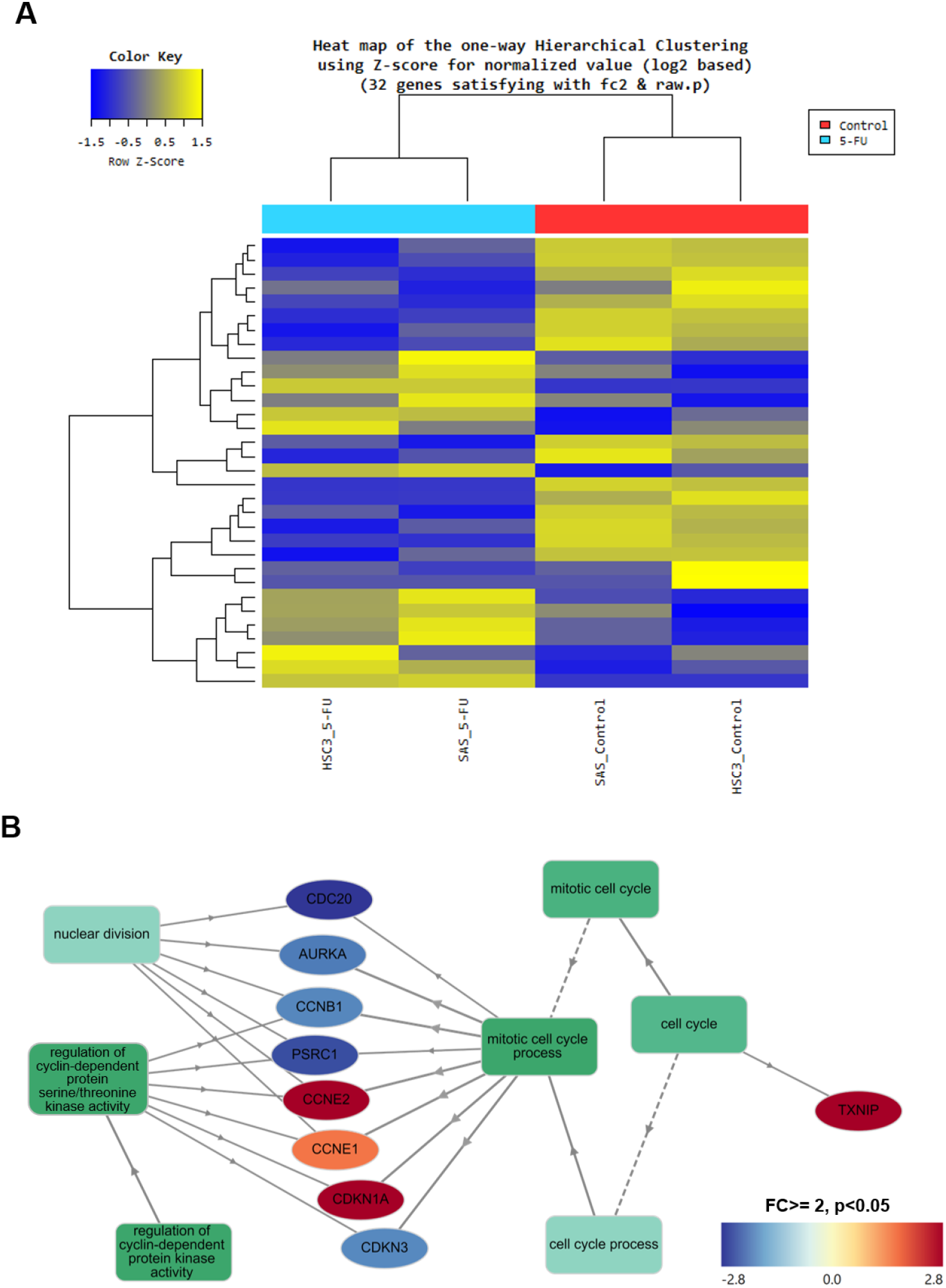
Transcriptome sequencing analysis. (A) Heat map shows result of hierarchical clustering analysis which clusters the similarity of genes and samples by expression level (normalized value) from a significant list. (B) GO analysis of significant expressed genes regarding cell cycle both in SAS and HSC3 after 10 μM 5-FU treatment for 16 h. Color density indicates the gene expression level.

**Figure 9.**
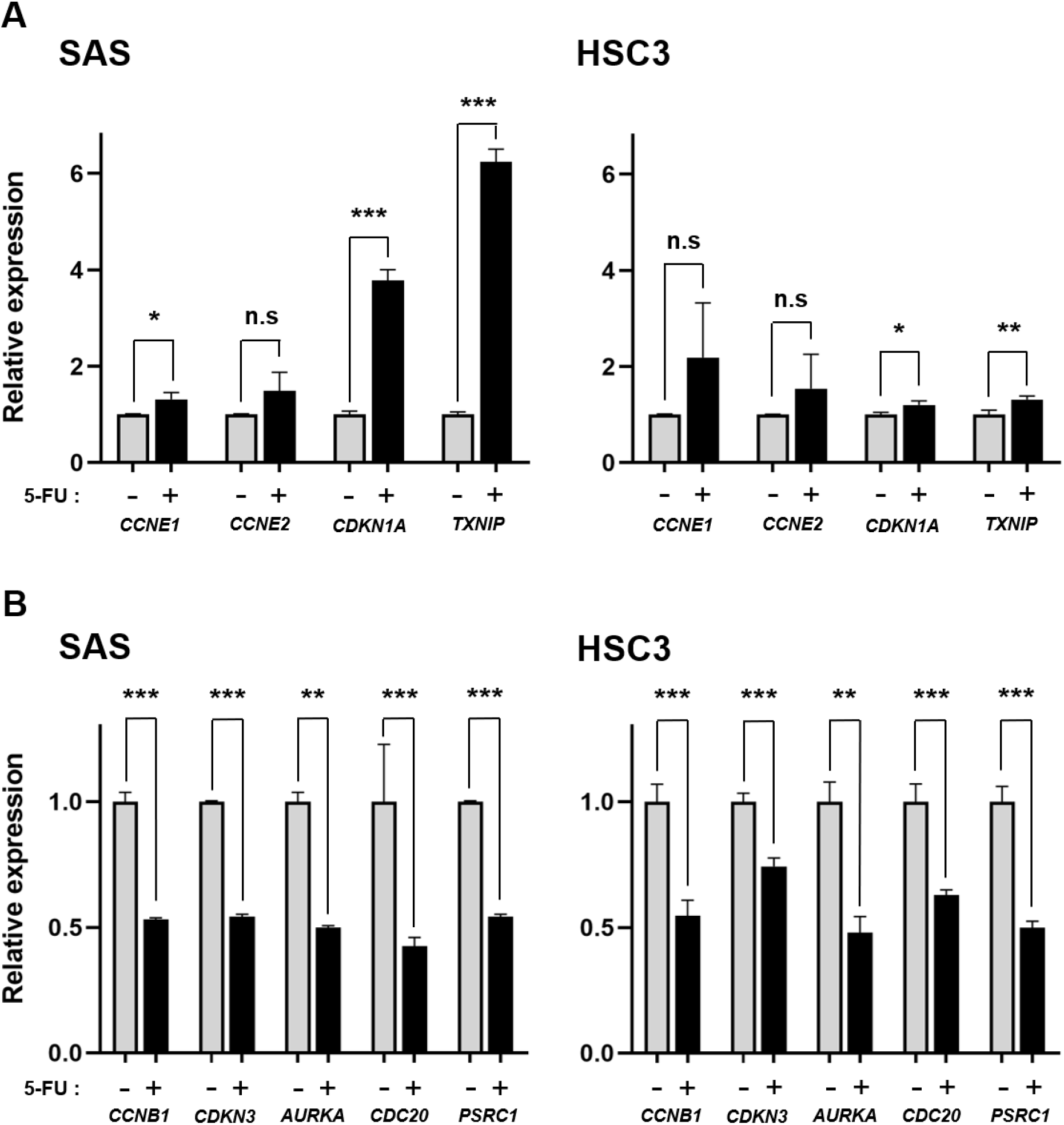
Gene expression analysis by qRT-PCR. (A, B) Quantitative qRT-PCR after 10 μM 5-FU treatment for 16 h in SAS and HSC3. All experiments were replicated 3 times. The values obtained were described as means ± SD. Data were compared statistically using the two-tailed Student’s *t*-test. *p-*value; *, ** and *** represent *p* < 0.05, *p* < 0.01 and *p* < 0.001, respectively.

### ATR inhibition enhanced efficacy of 5-FU independent of NHEJ and HR

5-FU treatment caused cell cycle arrest in S phase, where HR repair is active for repairing DSBs. BRCA2, one of the HR repair components, plays an important role in repairing DNA damage induced by 5-FU (3). To investigate whether ATR inhibition affects the sensitivity of 5-FU in the absence of BRCA2, surviving fractions of the Chinese hamster lung fibroblasts were measured. Contrary to our assumption that *BRCA2*-deficient cells would not be sensitized to ATRi treatment because BRCA2 is a downstream of ATR (25–28), we found that, compared with *BRCA2*-proficient cells, *BRCA2*-deficient cells were more sensitive to 5-FU in the presence of ATRi (Fig. 10A). This suggests that DNA damage induced by 5-FU treatment could also be repaired by mechanisms other than HR repair pathway.

**Figure 10.**
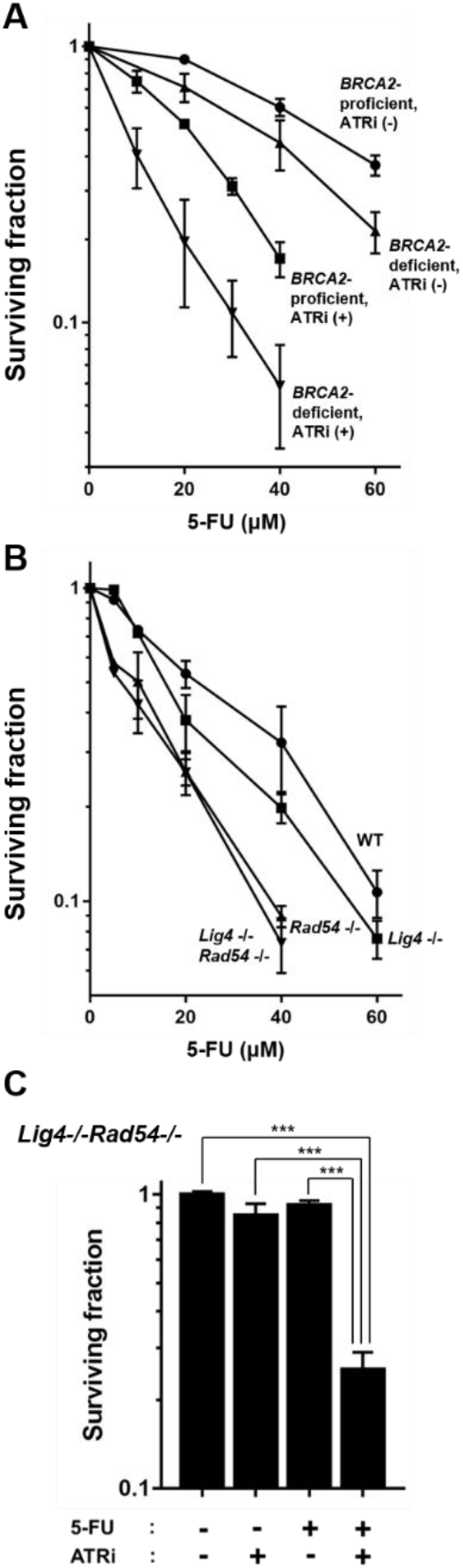
Contributions of NHEJ and HR repair pathways. (A) Surviving fraction of *BRCA2*- proficient and *BRCA2*-deficient Chinese hamster lung fibroblasts treated with various concentration of 5-FU and/or 3 μM ATRi for 24 h. (B) Surviving fraction of MEF WT (filled circles); *Lig4*−/− (filled squares); *Rad54*−/− (filled triangles); *Lig4*−/−*Rad54*−/− (filled inverted triangle) treated with 5-FU for 24 h. (C) Surviving fraction of MEF *Lig4*−/−*Rad54*−/− treated with 0.5 μM 5-FU and/or 0.5 μM ATRi for 24 h. All experiments were replicated 3 times. The values obtained were described as means ± SD. Data were compared statistically using the two-tailed Student’s *t*-test. *p-*value; *, ** and *** represent *p* < 0.05, *p* < 0.01 and *p* < 0.001, respectively.

To further determine whether the NHEJ or HR repair pathway is more predominant against 5-FU treatment, the surviving fraction of mouse embryonic fibroblasts (MEF) (WT; *Lig4*−/−; *Rad54*−/−; *Lig4*−/−*Rad54*−/−) was examined. All repair gene defective cells were sensitive to 5-FU but *Rad54*−/− cells were more sensitive to 5-FU than *Lig4*−/− cells. There was very little difference in the surviving fraction between *Rad54*−/− cells and *Lig4*−/−*Rad54*−/− cells (Fig. 10B). These results suggest that HR is more crucial than NHEJ in repairing DNA damage induced by 5-FU. Subsequently, we examined the effect of 5-FU treatment combined with ATRi to the *Lig4*−/− *Rad54*−/− cell lines. Surviving fractions of 0.5 μM ATRi alone, 0.5 μM 5-FU alone, and 0.5 μM 5-FU combined with 0.5 μM ATRi for 24 h was 85.0 ± 8.6%, 91.8 ± 3.5%, and 25.3 ± 3.7% respectively (Fig. 10C). Cell viability was remarkably decreased in the 5-FU treatment combined with ATRi compared to 5-FU alone (Fig.10C). In this case, neither HR nor NHEJ was involved in the repair of 5-FU induced DNA lesion. It is conceivable that SSB repair pathway like BER and MMR are responsible for the repair of 5-FU-induced DNA damage in the absence of both NHEJ and NR repair pathways. The significant difference of cell survival between 5-FU treatment alone and 5-FU combined with ATRi might not only be due to ATRi suppression of SSB repair but also its impact on molecular mechanisms other than DNA repair.

## Discussion

ATR is the principal protein for the repair of stalled replication forks (33). ATR senses stalled replication forks and is recruited to the forks through direct interactions with the ssDNA coated RPA at the forks, consequently preventing fork collapse and the formation of DNA breaks (34–36). Moreover, ATR is involved in S and G2 phase arrest by activating intra-S and G2/M checkpoint (33), and is necessary during HR repair pathway (37). On the other hand, 5-FU is thought to be an inhibitor of the enzyme TS, which plays a role in nucleotide synthesis (38,39). 5-FU induces unstable conformations in the DNA structure at the S phase, and where too many SSBs are present at stalled replication forks in 5-FU-treated cells, DSBs are induced (9,10). We consider ATR to be the principal factor in recognizing and repairing DNA damage induced by 5-FU treatment.

To repair DNA damage accurately, cell cycle must be arrested by cell cycle checkpoint to allow time for DNA repair. It was previously reported that 5-FU treatment led to S phase arrest (9) and our results in this study showed that cells were arrested at S phase after 5-FU treatment (Fig. 7A). ATR activates intra-S checkpoint in response to DNA damage (33), and inhibition of ATR suppresses the intra-S checkpoint leading the cells with DNA damage to enter G2 phase (40). Subsequently, damaged cells at G2 phase enter mitosis by the effect of ATRi (40), and then mitotic catastrophe occurs during mitosis (41,42). In the presence of ATRi, cells are not capable of activating intra-S checkpoint and repairing DNA damage induced by 5-FU. Thus, cell death induced by combination of 5-FU and ATRi might be caused by mitotic catastrophe.

Based on the results of the transcriptome analysis together with qPCR, 5-FU treatment led to cell cycle arrest between S phase and mitotic phase, especially at S phase (Fig. 8B, 9A and B). CCNE (CCNE1 and CCNE2), encoded by *CCNE1* and *CCNE2*, are involved in G1/S transition, and its expression level gradually increases as the cell cycle transitions from G1 phase to S phase, reaching its highest expression level immediately after entering S phase and gradually being degraded through S phase (43). Cyclin-dependent kinase (CDK) inhibitor p21 encoded by *CDKN1A* arrests the cell cycle at G1, S and G2 phases by preventing cyclin-CDK complex (44–46). TXNIP encoded by *TXNIP* acts on cell cycle arrest through retaining p27/CDK inhibitor in the nucleus (47,48). High expressions of *CCNE1* and *CCNE2* in response to 5-FU treatment suggests that 5-FU treatment causes early S phase arrest, and high expression of *CDKN1A* and *TXNIP* after 5-FU treatment explains why cells were arrested in S phase after 5-FU treatment (Fig. 7A). CCNB1 encoded by *CCNB1* is involved in G2/M transition (49), and expression of CCNB1 increases as the G2 phase progresses (50), thus *CCNB1* downregulation suggests that cells were arrested at G2 phase in response to 5-FU treatment. Proteins encoded by *CDC20*, *AURKA* and *PSRC1* play roles in mitosis (51–53), and the downregulation of these three genes suggests that cells were arrested at mitotic phase in response to 5-FU treatment.

ATR was reported to be recruited to centromeres in mitosis dependent on the activity of Aurora A kinase encoded by *AURKA* (54). The downregulation of *AURKA* in response to 5-FU treatment suggests that 5-FU treatment could inhibit the localization of ATR to centromeres. ATR inhibition on top of 5-FU treatment might further compromise centromere maintenance, leading to increased cell death at mitotic phase.

The auto-phosphorylation of ATM, an indicator of ATM activation, was induced by 5-FU treatment (Fig. 1). However, in terms of cell killing, ATMi was not effective as ATRi when combined with 5-FU (Fig. 2A, B and D). *CDKN3* encodes the downstream effector of ATM in DSBs repair pathway, which is a KRAB associated protein (KAP-1), and KAP-1 is phosphorylated at DSBs damage site in ATM dependent manner (55). *CDKN3* was downregulated in response to 5-FU, suggesting that low amount of KAP-1 leads to deficit in the repair of DSBs conducted by ATM even though ATM was activated by 5-FU treatment.

ATR inhibition using *BRCA2*-deficient cells and *Lig4*/*Rad54*-knockout cells demonstrates that ATR is involved in DNA repair other than NHEJ and HR (Fig. 10A, B and C). ATR responds to a wide range of DNA damage and DNA replication stress (56), and it is required for telomere maintenance through alternative lengthening of telomeres (57). In addition, ATR plays key roles in the suppression of chromosome instability at centromere through the promotion of faithful chromosome segregation (54). 5-FU treatment combined with ATRi is effective on cell killing because ATR inhibition not only blocks DNA repair pathways but also affects other intracellular dynamics, such as chromosome maintenance through telomeres and centromeres.

The results in our current study suggest that ATR inhibition is a potential therapeutic approach to enhance 5-FU treatment on cancer cells. With different *p53*-status, they are not only dependent on the HR repair pathway, but also other DNA repair pathways. For future therapeutic efforts, the application of ATR inhibitor may prove to be an effective tool in enhancing the efficacy of 5-FU chemotherapy for cancer patients. Subsequent studies will be required to further elucidate mechanism of DDR and the repair of 5-FU-induced DNA damage.

## Experimental procedures

### Cell lines

The present study used SAS (*p53*-proficient) and HSC3 (*p53*-deficient) human oral squamous cell carcinoma cell lines obtained from the Japanese Collection of Research Bioresources (Health Science Research Resources Bank, Osaka, Japan). SAS cells express wild-type p53 protein (58–60). HSC3 cells are impaired to express p53 protein (61). Chinese hamster lung fibroblasts used were V79 (*BRCA2*-proficient) and V-C8 (*BRCA2*-deficient) kindly provided by Dr. M.Z. Zdzienicka. The cell lines used were mouse embryonic fibroblasts (MEF) *Lig4*+/+*Rad54*+/+*p53*−/− (*WT*); *Lig4*−/− *Rad54*+/+*p53*−/− (*Lig4*−/−); *Lig4*+/+*Rad54*−/−*p53*−/− (*Rad54*−/−); *Lig4*−/−*Rad54*−/−*p53*−/− (*Lig4*−/−*Rad54*−/−) kindly provided by Dr. F.W. Alt. Cells were cultured at 37°C in Dulbecco’s modified Eagle’s medium containing 10% (v/v) fetal bovine serum, penicillin (50 U/ml), streptomycin (50 μg/ml) (DMEM-10).

### Chemicals and chemical treatment

5-FU (Kyowa Hakko, Tokyo, Japan); ATR inhibitor VE-821 (Selleck chemicals, Houston, USA); DNA-PK inhibitor NU7441 (KU57788) (TOCRIS, Bristol, UK); and ATM inhibitor KU55933 (TOCRIS, Bristol, UK) were used either on its own or in combination. A medium containing 5-FU at various concentrations was used to treat cells with respective inhibitors over a range of 8 h to 24 h before the cells were rinsed twice with PBS.

### Western blotting

Total protein from SAS cells treated with 5-FU and/or ATRi for 24 h were isolated using RIPA lysis buffer containing protease and phosphatase inhibitor cocktail, and quantified by the Protein Assay Bicinchoninate Acid (BCA) Kit (Nacalai Tesque, Kyoto, Japan). Isolated proteins (30 μg) were separated by 4-15% sodium dodecyl sulfate-polyacrylamide gel electrophoresis (SDS-PAGE) and transferred onto PVDF membranes. The membranes were probed overnight at 4°C using the following antibodies: anti-phospho-ATR (Thr-1989; GTX128145, GeneTex, Los Angeles, USA), anti-ATR (A300-138A, Bethyl, Montgomery, USA), anti-phospho-ATM (Ser-1981; #13050, Cell Signaling Technology, Danvers, USA), anti-ATM (A300-299A, Bethyl, Montgomery, USA), anti-phospho-DNA-PKcs (Ser-2056; ab124918, Abcam, Cambridge, UK), anti-DNA-PKcs (3H6; #123111, Cell Signaling Technology, Danvers, USA), anti-phospho-Chk1 (Ser-345; #2341, Cell Signaling Technology, Danvers, USA), anti-Chk1 (G-4; SC-8408, Santa Cruz, Dallas, USA), anti-p53 (DO-1; SC-126, Santa Cruz, Dallas, USA) and anti-Beta-Actin (Wako, Osaka, Japan). Blots were visualized using an enhanced chemiluminescence method (Bio-Rad Laboratories, Hercules, USA) according to the manufacturer’s protocol.

### Colony forming assays

We measured cell survival using a standard colony forming assay. In each experiment, three flasks were used, and three independent experiments were performed at each survival point. Colonies obtained after 7-10 days were fixed with methanol and stained with a 2% Giemsa solution. Microscopic colonies containing approximately 50 cells were scored as having grown from single surviving cells.

### DSBs analysis by neutral comet assay

Neutral single cell gel electrophoresis (comet) assay was performed using Comet Assay Kit (CELL BIOLABS, San Diego, USA). The treated cells were re-suspended at 10^5^ cells/ml in ice-cold PBS. Combined cell samples with Comet Agarose at 1:10 ratio, mixed well by pipetting, were immediately transferred to 75 μl/well. The slide was then transferred to a pre-chilled Lysis Buffer for 30 min, and then transferred to a pre-chilled Alkaline Solution for 30 min. It was subsequently immersed in pre-chilled neural TBE Electrophoresis Solution. Voltage was applied to the immersed slide for 15 min at 1 volt/cm. After electrophoresis, the slide was stained with 1:10,000 diluted Vista Green DNA Dye. Nuclei were observed under a fluorescence microscope. Each comet tail moment was quantified using ImageJ (62).

### H2AX phosphorylation analysis by immunocytochemistry

Cells were grown on glass slides in 6-well plates, fixed in 2% PFA in PBS for 15 minutes at room temperature. We permeabilized the cells for 5 minutes at 4°C in 0.2% Triton X-100, and they were blocked in PBS with 1% Bovine Serum Albumin (BSA) for 1 h at 37°C. They were then incubated with anti-phospho-H2AX (Ser-139) mouse monoclonal antibody (Upstate Biotechnology, Lake Placid, USA) for 1 h at 1:300 dilutions in PBS containing 1% BSA, and washed three times in PBS containing 1% BSA for 10 minutes. The cells were incubated with AlexaFluor 488-conjugated anti-mouse second antibody (Molecular Probes, Eugene, USA) for 1 h at room temperature at 1:400 dilutions in PBS containing 1% BSA, and washed three times for 10 minutes in PBS. Cover-glasses were mounted at 1:1000 dilutions of 4’,6-diamidino-2-phenylindole. Fluorescent images were captured for analysis using a fluorescence microscope.

### H2AX phosphorylation analysis by flow cytometry

Cells were fixed in cold 70% methanol after a 10 μM 5-FU with or without ATRi (3 μM) treatment for 6 h and 12 h, and maintained at 4°C up to 1 week before analysis. The overall levels of phosphorylated H2AX (γH2AX) were measured with flow cytometry.

### Analysis of apoptosis by Hoechst staining

Detection of apoptotic bodies with a Hoechst33342 staining assay was used to analyze the induction of apoptosis. Cells were fixed with 1% glutaraldehyde (Nacalai Tesque, Kyoto Japan) in PBS at 4°C, washed with PBS, stained with 0.2mM Hoechst33342 (Nacalai Tesque, Kyoto Japan) and then observed under a fluorescence microscope.

### Analysis of apoptosis by flow cytometry

After 5-FU and/or ATRi treatment, cells were fixed with cold 70% methanol and stored at 4°C for three days prior to analysis. To analyze the cell cycle, cells were incubated for 30 minutes at room temperature with 1 mg/mL RNase and 50 μg/mL propidium iodoide (PI), and analyzed using a flow cytometer. Cell cycle distribution was then assayed by determining the DNA content twice and deriving its average values.

### Transcriptome sequencing analysis

Total RNA from SAS and HSC3 cells was isolated according to the protocol specified in the Purelink RNA mini kit (Thermo Fisher Scientific, Waltham, USA). We pooled three replicated samples of each RNA into one sample. We used DU730 UV-vis Spectrophotometer (Beckman Coulter, Brea, USA) to measure the concentration and purity of the RNA samples. Contamination DNA was eliminated using DNase. RNA was purified randomly fragment for short read sequencing, and then reverse transcribed into cDNA. Adapters were ligated onto both ends of the cDNA fragments. After amplifying fragments using PCR, selected fragments with insert sizes between 200-400 bp. For paired-end sequencing, both ends of the cDNA was sequenced by the read length. The quality control of the sequenced raw reads was analyzed. Trimmed reads were mapped to reference genome with HISAT2 (63,64), splice-aware aligner. Transcript was assembled by StringTie (64,65) with aligned reads. Expression profiles were represented as read count and normalization value which is based on transcript length and depth of coverage. In groups with different conditions, genes or transcripts that express differentially were filtered out though statistical hypothesis testing. Statistical analysis was performed using fold change, exactTest (66) using edgeR (67) per comparison pair. The significant results were selected on conditions of |fc| >= 2 and raw *p*-value < 0,05. In case of known gene annotation, functional annotation and gene-enrichment analysis were performed using GO net (68) based on GO (http://geneontology.org/) database.

### Quantitative PCR

RNA from SAS and HSC3 cells was extracted according to the protocol specified in the Purelink RNA mini kit (Thermo Fisher Scientific, Waltham, USA). We used DU730 UV-vis Spectrophotometer (Beckman Coulter, Brea, USA) to measure the concentration and purity of the RNA samples. Extracted RNA (0.5 μg) was reverse-transcribed based on the protocol outlined in the ReverTra Ace qPCR RT Master Mix (TOYOBO, Osaka, Japan). StepOne Plus Real-time PCR System (Thermo Fisher Scientific, Waltham, USA) was used to amplify and quantify levels of target gene cDNA. We performed quantitative real-time PCR (qRT-PCR) with SsoAdvanced Universal SYBR Green Supermix (Bio-Rad Laboratories, Hercules, USA) and specific primers for qRT-PCR. We also ran reactions in triplicate and normalized the expression of each gene to the geo-metric mean of β-actin as a housekeeping gene, and applied the ΔΔCT method for analysis. Following primers were designed by Bio-Rad; CCNE1 (qHsaCID0015131), CCNE2 (qHsaCID0007224), CCNB1 (qHsaCED0044529), CDKN1A (qHsaCID0014498), CDKN3 (qHsaCID0010035), CDC20 (qHsaCID0012637), AURKA (qHsaCID0022123), TXNIP (qHsaCED0043730), PSRC1 (qHsaCED0038868), β-actin (qHsaCED0036269).

### Data availability statement

The bulk RNA sequencing reads have been submitted to the DDBJ Sequence Read Archive (DRA) under the accession number SAMD00220269 (HSC3 5-FU), SAMD00220270 (HSC3 Control), SAMD00220271 (SAS 5-FU) and SAMD00220272 (SAS Control).

### Statistical analysis

The values obtained were described as means ± standard deviations (SD). Data were compared statistically using the two-tailed Student’s *t*-test. *p-*value; */†, **/†† and ***/††† represent *p* < 0.05, *p* < 0.01 and *p* < 0.001, respectively.

## Conflict of interest

The authors declare that they have no conflicts of interest with the contents of article.

## Acknowledgements

The authors thank Dr. Takeo Ohnishi for his guidance of and support for this project over these years, Keren-Happuch E and Dr. Takahiko Nakagawa for their critical reading of the manuscript, and Drs. M.Z. Zdzienicka and F.W. Alt. for kindly providing the cell lines used in this work. This work was supported by Takeda Science Foundation to E.M. and T.K.M., Kanzawa Medical Research Foundation to E.M., Uehara Memorial Foundation to E.M. and S.Kikuchi., Nakatomi Foundation to E.M., Konica Minolta Science and Technology Foundation to E.M., Naito Foundation to E.M., MSD Life Science Foundation to E.M., Mochida Memorial Foundation for Medical and Pharmaceutical Research to E.M., SENSHIN Medical Research Foundation to E.M., Terumo Foundation for Life Sciences and Arts to E.M., Nara Kidney Disease Research Foundation to E.M., Novartis Research Grants to E.M. and H.N., Sumitomo Dainippon Pharma Research Grant to T.K.M., Tokyo Biochemical Research Foundation to S.Kikuchi. and the unrestricted funds provided to E.M. from Dr. Taichi Noda (KTX Corp., Aichi, Japan) and Dr. Yasuhiro Horii (Koseikan, Nara, Japan).

## Footnotes

Funding was provided by grants from JSPS KAKENHI [JP18K17234 to Y.N., JP20H03199 to E.M., JP19K08150 to S.Kobashigawa., JP19K23976 to M.N., JP19K21306 and JP20K16583 to H.N., JP18K07764 to G.K., and JP19K10272 to T.K.] and AMED Brain/MINDS Beyond [JP20dm0307032 to E.M.]

The abbreviations used are:

ATR: ataxia telangiectasia mutated and Rad3-related protein
5-FU: 5-fluorouracil
DSBs: double-strand breaks
NHEJ: non-homologous end joining
HR: homologous recombination
SSBs: single-strand breaks
ssDNA: single-strand DNA
ATM: ataxia telangiectasia mutated protein
ATRIP: ATR interacting protein
DNA-PKcs: DNA-dependent protein kinase catalytic subunit
Chk1: checkpoint kinase 1
BRCA2: breast cancer susceptibility gene 2
CCNB1: Cyclin B1
CCNE1: Cyclin E1
CCNE2: Cyclin E2
CDKN1A: Cyclin dependent kinase inhibitor 1A
CDKN3: Cyclin dependent kinase inhibitor 3
CDC20: Cell division cycle 20
AURKA: Aurora kinase A
TXNIP: Thioredoxin interacting protein
PSRC1: Proline and serine rich coiled-coil 1
MEF: mouse embryonic fibroblasts
KAP-1: KRAB associated protein
logCPM: log counts per million

## Reference

1. Longley, D. B., Harkin, D. P., and Johnston, P. G. (2003) 5-fluorouracil: mechanisms of action and clinical strategies. Nat Rev Cancer 3, 330–338

2. Miura, K., Kinouchi, M., Ishida, K., Fujibuchi, W., Naitoh, T., Ogawa, H., Ando, T., Yazaki, N., Watanabe, K., Haneda, S., Shibata, C., and Sasaki, I. (2010) 5-fu metabolism in cancer and orally-administrable 5-fu drugs. Cancers (Basel) 2, 1717–1730

3. Nakagawa, Y., Kajihara, A., Takahashi, A., Kondo, N., Mori, E., Kirita, T., and Ohnishi, T. (2014) The BRCA2 gene is a potential molecular target during 5-fluorouracil therapy in human oral cancer cells. Oncol Rep 31, 2001–2006

4. Wyatt, M. D., and Wilson, D. M. (2009) Participation of DNA repair in the response to 5-fluorouracil. Cell Mol Life Sci 66, 788–799

5. Li, L. S., Morales, J. C., Veigl, M., Sedwick, D., Greer, S., Meyers, M., Wagner, M., Fishel, R., and Boothman, D. A. (2009) DNA mismatch repair (MMR)-dependent 5-fluorouracil cytotoxicity and the potential for new therapeutic targets. Br J Pharmacol 158, 679–692

6. Fischer, F., Baerenfaller, K., and Jiricny, J. (2007) 5-Fluorouracil is efficiently removed from DNA by the base excision and mismatch repair systems. Gastroenterology 133, 1858–1868

7. Das, D., Preet, R., Mohapatra, P., Satapathy, S. R., Siddharth, S., Tamir, T., Jain, V., Bharatam, P. V., Wyatt, M. D., and Kundu, C. N. (2014) 5-Fluorouracil mediated anti-cancer activity in colon cancer cells is through the induction of Adenomatous Polyposis Coli: Implication of the long-patch base excision repair pathway. DNA Repair (Amst) 24, 15–25

8. Lindahl, T. (1974) An N-glycosidase from Escherichia coli that releases free uracil from DNA containing deaminated cytosine residues. Proc Natl Acad Sci U S A 71, 3649–3653

9. Fujinaka, Y., Matsuoka, K., Iimori, M., Tuul, M., Sakasai, R., Yoshinaga, K., Saeki, H., Morita, M., Kakeji, Y., Gillespie, D. A., Yamamoto, K., Takata, M., Kitao, H., and Maehara, Y. (2012) ATR-Chk1 signaling pathway and homologous recombinational repair protect cells from 5-fluorouracil cytotoxicity. DNA Repair (Amst) 11, 247–258

10. Zhang, X. P., Liu, F., Cheng, Z., and Wang, W. (2009) Cell fate decision mediated by p53 pulses. Proc Natl Acad Sci U S A 106, 12245–12250

11. Maréchal, A., and Zou, L. (2013) DNA damage sensing by the ATM and ATR kinases. Cold Spring Harb Perspect Biol 5

12. Woods, D., and Turchi, J. J. (2013) Chemotherapy induced DNA damage response: convergence of drugs and pathways. Cancer Biol Ther 14, 379–389

13. Heffernan, T. P., Simpson, D. A., Frank, A. R., Heinloth, A. N., Paules, R. S., Cordeiro-Stone, M., and Kaufmann, W. K. (2002) An ATR- and Chk1-dependent S checkpoint inhibits replicon initiation following UVC-induced DNA damage. Mol Cell Biol 22, 8552–8561

14. Borges, H. L., Linden, R., and Wang, J. Y. (2008) DNA damage-induced cell death: lessons from the central nervous system. Cell Res 18, 17–26

15. Goto, H., Natsume, T., Kanemaki, M. T., Kaito, A., Wang, S., Gabazza, E. C., Inagaki, M., and Mizoguchi, A. (2019) Chk1-mediated Cdc25A degradation as a critical mechanism for normal cell cycle progression. J Cell Sci 132

16. Buisson, R., Boisvert, J. L., Benes, C. H., and Zou, L. (2015) Distinct but Concerted Roles of ATR, DNA-PK, and Chk1 in Countering Replication Stress during S Phase. Mol Cell 59, 1011–1024

17. Jackson, S. P. (2002) Sensing and repairing DNA double-strand breaks. Carcinogenesis 23, 687–696

18. Kanaar, R., Hoeijmakers, J. H., and van Gent, D. C. (1998) Molecular mechanisms of DNA double strand break repair. Trends Cell Biol 8, 483–489

19. Rothkamm, K., Krüger, I., Thompson, L. H., and Löbrich, M. (2003) Pathways of DNA double-strand break repair during the mammalian cell cycle. Mol Cell Biol 23, 5706–5715

20. Hinz, J. M., Yamada, N. A., Salazar, E. P., Tebbs, R. S., and Thompson, L. H. (2005) Influence of double-strand-break repair pathways on radiosensitivity throughout the cell cycle in CHO cells. DNA Repair (Amst) 4, 782–792

21. Thompson, L. H., and Schild, D. (2001) Homologous recombinational repair of DNA ensures mammalian chromosome stability. Mutat Res 477, 131–153

22. Davies, A. A., Masson, J. Y., McIlwraith, M. J., Stasiak, A. Z., Stasiak, A., Venkitaraman, A. R., and West, S. C. (2001) Role of BRCA2 in control of the RAD51 recombination and DNA repair protein. Mol Cell 7, 273–282

23. Ottini, L., Masala, G., D’Amico, C., Mancini, B., Saieva, C., Aceto, G., Gestri, D., Vezzosi, V., Falchetti, M., De Marco, M., Paglierani, M., Cama, A., Bianchi, S., Mariani-Costantini, R., and Palli, D. (2003) BRCA1 and BRCA2 mutation status and tumor characteristics in male breast cancer: a population-based study in Italy. Cancer Res 63, 342–347

24. Schrader, K. A., Hurlburt, J., Kalloger, S. E., Hansford, S., Young, S., Huntsman, D. G., Gilks, C. B., and McAlpine, J. N. (2012) Germline BRCA1 and BRCA2 mutations in ovarian cancer: utility of a histology-based referral strategy. Obstet Gynecol 120, 235–240

25. Matsuoka, S., Ballif, B. A., Smogorzewska, A., McDonald, E. R., Hurov, K. E., Luo, J., Bakalarski, C. E., Zhao, Z., Solimini, N., Lerenthal, Y., Shiloh, Y., Gygi, S. P., and Elledge, S. J. (2007) ATM and ATR substrate analysis reveals extensive protein networks responsive to DNA damage. Science 316, 1160–1166

26. Gatei, M., Zhou, B. B., Hobson, K., Scott, S., Young, D., and Khanna, K. K. (2001) Ataxia telangiectasia mutated (ATM) kinase and ATM and Rad3 related kinase mediate phosphorylation of Brca1 at distinct and overlapping sites. In vivo assessment using phospho-specific antibodies. J Biol Chem 276, 17276–17280

27. Fradet-Turcotte, A., Sitz, J., Grapton, D., and Orthwein, A. (2016) BRCA2 functions: from DNA repair to replication fork stabilization. Endocr Relat Cancer 23, T1–T17

28. Venkitaraman, A. R. (2001) Functions of BRCA1 and BRCA2 in the biological response to DNA damage. J Cell Sci 114, 3591–3598

29. Liu, S., Shiotani, B., Lahiri, M., Maréchal, A., Tse, A., Leung, C. C., Glover, J. N., Yang, X. H., and Zou, L. (2011) ATR autophosphorylation as a molecular switch for checkpoint activation. Mol Cell 43, 192–202

30. Takahashi, A., and Ohnishi, T. (2005) Does gammaH2AX foci formation depend on the presence of DNA double strand breaks? Cancer Lett 229, 171–179

31. Darzynkiewicz, Z., Bruno, S., Del Bino, G., Gorczyca, W., Hotz, M. A., Lassota, P., and Traganos, F. (1992) Features of apoptotic cells measured by flow cytometry. Cytometry 13, 795–808

32. Bedner, E., Li, X., Gorczyca, W., Melamed, M. R., and Darzynkiewicz, Z. (1999) Analysis of apoptosis by laser scanning cytometry. Cytometry 35, 181–195

33. Saldivar, J. C., Cortez, D., and Cimprich, K. A. (2017) The essential kinase ATR: ensuring faithful duplication of a challenging genome. Nat Rev Mol Cell Biol 18, 622–636

34. Smith, J., Tho, L. M., Xu, N., and Gillespie, D. A. (2010) The ATM-Chk2 and ATR-Chk1 pathways in DNA damage signaling and cancer. Adv Cancer Res 108, 73–112

35. Feijoo, C., Hall-Jackson, C., Wu, R., Jenkins, D., Leitch, J., Gilbert, D. M., and Smythe, C. (2001) Activation of mammalian Chk1 during DNA replication arrest: a role for Chk1 in the intra-S phase checkpoint monitoring replication origin firing. J Cell Biol 154, 913–923

36. Cortez, D. (2015) Preventing replication fork collapse to maintain genome integrity. DNA Repair (Amst) 32, 149–157

37. Sirbu, B. M., and Cortez, D. (2013) DNA damage response: three levels of DNA repair regulation. Cold Spring Harb Perspect Biol 5, a012724

38. Welsh, S. J., Hobbs, S., and Aherne, G. W. (2003) Expression of uracil DNA glycosylase (UDG) does not affect cellular sensitivity to thymidylate synthase (TS) inhibition. Eur J Cancer 39, 378–387

39. Koehler, S. E., and Ladner, R. D. (2004) Small interfering RNA-mediated suppression of dUTPase sensitizes cancer cell lines to thymidylate synthase inhibition. Mol Pharmacol 66, 620–626

40. Qiu, Z., Oleinick, N. L., and Zhang, J. (2018) ATR/CHK1 inhibitors and cancer therapy. Radiother Oncol 126, 450–464

41. Mc Gee, M. M. (2015) Targeting the Mitotic Catastrophe Signaling Pathway in Cancer. Mediators Inflamm 2015, 146282

42. Nitta, M., Kobayashi, O., Honda, S., Hirota, T., Kuninaka, S., Marumoto, T., Ushio, Y., and Saya, H. (2004) Spindle checkpoint function is required for mitotic catastrophe induced by DNA-damaging agents. Oncogene 23, 6548–6558

43. Clurman, B. E., Sheaff, R. J., Thress, K., Groudine, M., and Roberts, J. M. (1996) Turnover of cyclin E by the ubiquitin-proteasome pathway is regulated by cdk2 binding and cyclin phosphorylation. Genes Dev 10, 1979–1990

44. Niculescu, A. B., Chen, X., Smeets, M., Hengst, L., Prives, C., and Reed, S. I. (1998) Effects of p21(Cip1/Waf1) at both the G1/S and the G2/M cell cycle transitions: pRb is a critical determinant in blocking DNA replication and in preventing endoreduplication. Mol Cell Biol 18, 629–643

45. Ogryzko, V. V., Wong, P., and Howard, B. H. (1997) WAF1 retards S-phase progression primarily by inhibition of cyclin-dependent kinases. Mol Cell Biol 17, 4877–4882

46. Radhakrishnan, S. K., Feliciano, C. S., Najmabadi, F., Haegebarth, A., Kandel, E. S., Tyner, A. L., and Gartel, A. L. (2004) Constitutive expression of E2F-1 leads to p21-dependent cell cycle arrest in S phase of the cell cycle. Oncogene 23, 4173–4176

47. Russo, A. A., Jeffrey, P. D., Patten, A. K., Massagué, J., and Pavletich, N. P. (1996) Crystal structure of the p27Kip1 cyclin-dependent-kinase inhibitor bound to the cyclin A-Cdk2 complex. Nature 382, 325–331

48. Yamaguchi, F., Takata, M., Kamitori, K., Nonaka, M., Dong, Y., Sui, L., and Tokuda, M. (2008) Rare sugar D-allose induces specific up-regulation of TXNIP and subsequent G1 cell cycle arrest in hepatocellular carcinoma cells by stabilization of p27kip1. Int J Oncol 32, 377–385

49. Petri, E. T., Errico, A., Escobedo, L., Hunt, T., and Basavappa, R. (2007) The crystal structure of human cyclin B. Cell Cycle 6, 1342–1349

50. Pines, J., and Hunter, T. (1989) Isolation of a human cyclin cDNA: evidence for cyclin mRNA and protein regulation in the cell cycle and for interaction with p34cdc2. Cell 58, 833–846

51. Fang, G., Yu, H., and Kirschner, M. W. (1999) Control of mitotic transitions by the anaphase-promoting complex. Philos Trans R Soc Lond B Biol Sci 354, 1583–1590

52. Nikonova, A. S., Astsaturov, I., Serebriiskii, I. G., Dunbrack, R. L., and Golemis, E. A. (2013) Aurora A kinase (AURKA) in normal and pathological cell division. Cell Mol Life Sci 70, 661–687

53. Jang, C. Y., Wong, J., Coppinger, J. A., Seki, A., Yates, J. R., and Fang, G. (2008) DDA3 recruits microtubule depolymerase Kif2a to spindle poles and controls spindle dynamics and mitotic chromosome movement. J Cell Biol 181, 255–267

54. Kabeche, L., Nguyen, H. D., Buisson, R., and Zou, L. (2018) A mitosis-specific and R loop-driven ATR pathway promotes faithful chromosome segregation. Science 359, 108–114

55. Ziv, Y., Bielopolski, D., Galanty, Y., Lukas, C., Taya, Y., Schultz, D. C., Lukas, J., Bekker-Jensen, S., Bartek, J., and Shiloh, Y. (2006) Chromatin relaxation in response to DNA double-strand breaks is modulated by a novel ATM- and KAP-1 dependent pathway. Nat Cell Biol 8, 870–876

56. Karnitz, L. M., and Zou, L. (2015) Molecular Pathways: Targeting ATR in Cancer Therapy. Clin Cancer Res 21, 4780–4785

57. Flynn, R. L., Cox, K. E., Jeitany, M., Wakimoto, H., Bryll, A. R., Ganem, N. J., Bersani, F., Pineda, J. R., Suvà, M. L., Benes, C. H., Haber, D. A., Boussin, F. D., and Zou, L. (2015) Alternative lengthening of telomeres renders cancer cells hypersensitive to ATR inhibitors. Science 347, 273–277

58. Abiko, Y., Arai, J., Mitamura, J., and Kaku, T. (1997) Alteration of proto-oncogenes during apoptosis in the oral squamous cell carcinoma cell line, SAS, induced by staurosporine. Cancer Lett 118, 101–107

59. Takahashi, A. (2001) Different inducibility of radiation- or heat-induced p53-dependent apoptosis after acute or chronic irradiation in human cultured squamous cell carcinoma cells. Int J Radiat Biol 77, 215–224

60. Kanata, H., Yane, K., Ota, I., Miyahara, H., Matsunaga, T., Takahashi, A., Ohnishi, K., Ohnishi, T., and Hosoi, H. (2000) CDDP induces p53-dependent apoptosis in tongue cancer cells. Int J Oncol 17, 513–517

61. Sakai, E., and Tsuchida, N. (1992) Most human squamous cell carcinomas in the oral cavity contain mutated p53 tumor-suppressor genes. Oncogene 7, 927–933

62. Schneider, C. A., Rasband, W. S., and Eliceiri, K. W. (2012) NIH Image to ImageJ: 25 years of image analysis. Nat Methods 9, 671–675

63. Kim, D., Langmead, B., and Salzberg, S. L. (2015) HISAT: a fast spliced aligner with low memory requirements. Nat Methods 12, 357–360

64. Pertea, M., Kim, D., Pertea, G. M., Leek, J. T., and Salzberg, S. L. (2016) Transcript-level expression analysis of RNA-seq experiments with HISAT, StringTie and Ballgown. Nat Protoc 11, 1650–1667

65. Pertea, M., Pertea, G. M., Antonescu, C. M., Chang, T. C., Mendell, J. T., and Salzberg, S. L. (2015) StringTie enables improved reconstruction of a transcriptome from RNA-seq reads. Nat Biotechnol 33, 290–295

66. Robinson, M. D., and Smyth, G. K. (2008) Small-sample estimation of negative binomial dispersion, with applications to SAGE data. Biostatistics 9, 321–332

67. Robinson, M. D., McCarthy, D. J., and Smyth, G. K. (2010) edgeR: a Bioconductor package for differential expression analysis of digital gene expression data. Bioinformatics 26, 139–140

68. Pomaznoy, M., Ha, B., and Peters, B. (2018) GOnet: a tool for interactive Gene Ontology analysis. BMC Bioinformatics 19, 470

